# Long-Term Neuromodulatory Effects on the Ionic Current Parameter Space of Oscillatory Neurons

**DOI:** 10.64898/2026.02.22.707287

**Authors:** Smita More-Potdar, Jorge Golowasch

## Abstract

Neurons are constantly subject to short-term neuromodulatory effects through the regulation of various proteins, including ion channels. However, the long-term influences of neuromodulators remain poorly understood. Multiple ionic currents work in concert to generate oscillatory neuronal activity, and long-term neuromodulatory effects on them likely differ from short-term effects. Here we examined the long-term effects of the neuromodulatory environment on the ionic conductances that generate the rhythmic activity of a well-known neuronal network, the crustacean pyloric network. We measured most of the known voltage-dependent and synaptic currents expressed by two identified neurons in their normal neuromodulatory environment and after prolonged removal of neuromodulators. To understand the global conductance makeup of these cells, we defined the conductance parameter space of each cell as the multidimensional volume occupied by its conductances in a population of identical neurons. We then examined the changes in this volume after independently modifying the neuromodulator environment and activity of these neurons. Both neuromodulation and activity appear to constrain the conductance parameter space in pyloric neurons, suggesting a general phenomenon. Interestingly, while neuromodulators appear to exert a similar regulatory effect on both cells, activity seems to do so in a cell-type-specific manner. Our results suggest a mechanism that explains why the same manipulation of the neuromodulatory input does not produce the same change in network activity across animals: the degeneracy of the pacemaker mechanism.

**Significance:** How does a rhythm-generating neuronal network recover its rhythm after its disruption? Neuromodulators control neuronal activity, especially rhythmic activity, which is vital to animals. Thus, neuronal network robustness and the ability to recover from major perturbations are of crucial importance. We tested the hypothesis that neuromodulators constrain the entire neuronal ionic conductance parameter space in a well-characterized network. Our results confirm that neuromodulators and activity regulate the global neuronal conductance landscape. Our results suggest a mechanism that explains why the same manipulation of the neuromodulatory environment produces distinct effects on network activity across individuals: the degeneracy of the pacemaker mechanism. These results support the need to consider individualized approaches to understand functional mechanisms and to develop therapies for pathological conditions.

## Introduction

The complex interplay among several voltage-dependent and synaptic currents is essential for the generation of neuronal oscillations that underlie many vital rhythmic activity patterns, *e*.*g*. feeding, locomotion, and breathing behaviors (Wei and Stengl, 2012; Ramirez and Baertsch, 2018; da Silva Junior et al., 2025). These activities must be reliably maintained throughout the lifetime of all animals to ensure their survival. This begs the question of how they are maintained in the face of potentially devastating changes due to disease, trauma, or large developmental changes. It is, therefore, crucial to understand how neurons coordinately regulate the full complement of their ionic currents rather than single or even small subsets of them, while maintaining stable activity. Central pattern generator networks (CPGs) often rely on neuromodulatory inputs to initiate, regulate, and fine-tune their rhythmic activity, typically acting GPCRs to influence ion channel function (Tryba et al., 2006; Golowasch, 2019a; Su et al., 2025). In the crab, *Cancer borealis*, the rhythmic movements of the food-filtering pyloric chamber are controlled by the pyloric CPG, which is regulated by a host of neuromodulators (Marder et al., 2022) (Fig. 1). Pyloric neurons express the modulator-activated inward current, *I*_MI_, a cationic current that is a direct and convergent target of several neuromodulators (Swensen and Marder, 2000; Schneider et al., 2021). It has been suggested that *I*_MI_ functions as a pacemaker current essential for generating rhythmic pyloric activity (Luther et al., 2003; Bose et al., 2014; Hamood et al., 2015; Gray et al., 2017) because deactivating *I*_MI_ typically stops oscillatory activity (Fig. 2A). However, it sometimes only slows rhythmic activity down to varying degrees (Fig. 2B). This suggests that *I*_MI_ activation is not the only pacemaking mechanism depending on the prior history of perturbations or modulatory conditions of the animal (Golowasch et al., 2002; O’Leary et al., 2014). Additionally, within hours of decentralization, the pyloric rhythm recovers many of its original activity features (Fig. 2A)without restoring its neuromodulatory inputs, and thus presumably *I*_MI_ (Thoby-Brisson and Simmers, 2000; Luther et al., 2003; More-Potdar and Golowasch, 2023). This recovery implies a compensatory reorganization at the level of all ion channels and also suggests the existence of multiple pacemaking mechanisms in the network.

**Figure 1.**
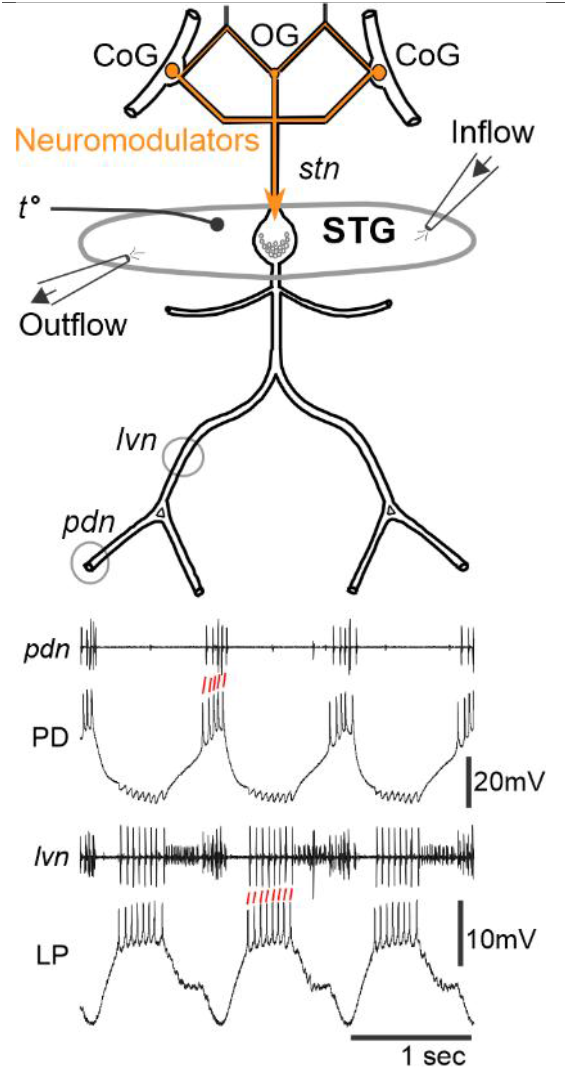
The Stomatogastric Nervous System (STNS). <u>Top</u>: Schematic diagram of the crab *Cancer borealis* STNS. CoG: Commissural Ganglion; OG: Oesophageal Ganglion; STG: Stomatogastric Ganglion; *stn*: stomatogastric nerve; *lvn*: lateral ventricular nerve; *pdn* pyloric dilator nerve (other nerve names have been omitted for clarity); *t*°: temperature probe. Oval around the STG is made of petroleum jelly (Vaseline). <u>Bottom</u>: Extracellular recordings obtained from two motor nerves of the pyloric system, the *pdn* and *lvn* nerves, with intracellular PD neuron recording under *pdn* and intracellular LP neuron recording under *lvn*. Red diagonal lines indicate matches between extracellular and intracellular action potentials used to identify the neurons. Note that the *pdn* recording has extra action potentials than those recorded intracellularly, which correspond to those generated by the second PD neuron in the ganglion (Fig. 2A). PD neurons release neurotransmitter in a slow, graded fashion (big dip in PD phase on LP neuron), while the LP neuron feeds back onto both PD neurons with fast inhibitory synaptic potentials.

**Figure 2.**
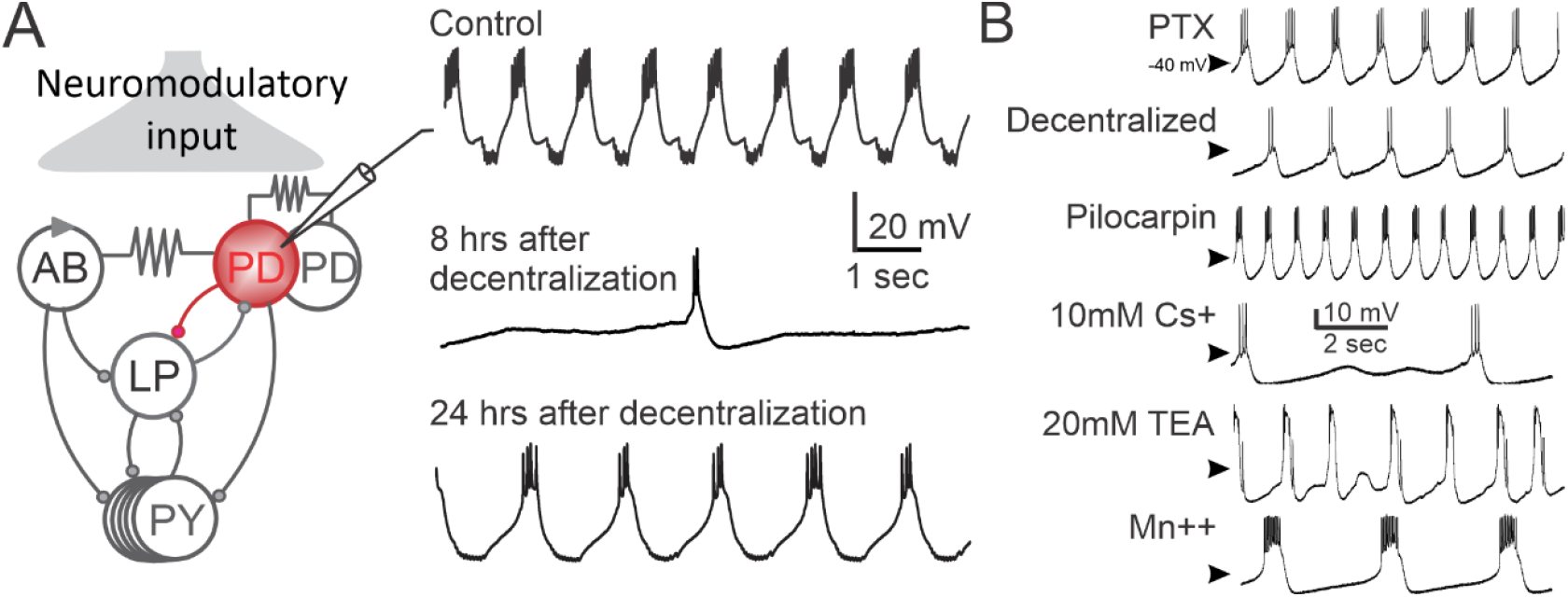
Pyloric network neurons generate regular oscillatory activity in the presence of neuromodulators. (**A**) Schematic illustration of the central part of the pyloric network, which includes the pacemaker kernel made of one Anterior Burster (AB) neuron and two PD neurons, and the single feedback neuron to the pacemaker, the LP neuron (left). Intracellular recordings from one PD neuron (right) show the pyloric rhythm in the presence of intact endogenous neuromodulatory input (Control),8 hours after the removal of neuromodulatory input (decentralization), and 24 hours after decentralization. The typical initial drastic slowing down of the pyloric rhythm and its almost complete recovery 24 hours later in the continued absence of neuromodulatory inputs are illustrated. All chemical synapses (circles are inhibitory; resistor represents electrical synapses. (**B**) Blocking individual ionic current in a PD neuron drastically affects its oscillatory features. Here, currents were blocked sequentially and were washed completely in between applications until control activity was fully restored (not shown). Initially, picrotoxin (‘PTX’) was used to remove all synaptic currents to the PD neuron (notice the absence of inhibitory potentials compared to Control in A) and was left in place through the following treatments: ‘Decentralized’ removes all neuromodulation and presumably *I*_MI_ (Golowasch and Marder, 1992a; Swensen and Marder, 2000); ‘Pilocarpine’, a muscarinic agonist, activates *I*_MI_ and restores a robust pyloric rhythm. From here on, pilocarpine was included in all treatments. ‘10mM Cs^+^’ blocks *I*_H_, ‘20mM TEA’ blocks *I*_KCa_ and partially also *I*_Kd_, and ‘Mn^++^’ blocks all Ca^++^ currents. Clearly, all ionic currents affect rhythmic activity, some slowing down the rhythm (Decentralization, Cs^+^), accelerating or activating it (Pilocarpine), and some changing its properties to different degrees (TEA, Mn^++^).

Neuromodulators exert well-documented short-term effects on their direct targets in various systems through intracellular signaling cascades (Gray et al., 2017; Park et al., 2023; Gonzalez-Hernandez et al., 2024), and some also have slower effects on indirect targets (Svensson et al., 2001; Garcia et al., 2015). It is also known that decentralization induces long-term changes in the expression of various voltage-gated and synaptic currents in pyloric neurons (Mizrahi et al., 2001; Thoby-Brisson and Simmers, 2002; Khorkova and Golowasch, 2007; Garcia et al., 2015), making them indirect targets of neuromodulators in pyloric neurons.

Whether they also regulate the entire conductance landscape (or parameter space) of each neuron to control rhythmic activity is not known. We hypothesize that the removal of neuromodulators (decentralization) not only modifies specific targets (*i*.*e. I*_MI_) in the short-term but also globally expands the conductance parameter space of pyloric neurons in the long-term, such that theseneurons find new solutions within this expanded parameter space, which may allow them to restore rhythmic activity (Golowasch, 2019a).

To understand the mechanisms that support the long-term maintenance of rhythmic function and its recovery after perturbations, we examined the largest possible ensemble of ionic currents in two identified neurons of the crab *C. borealis* pyloric network: one of the pacemaker neurons (Pyloric Dilator, PD neuron) and a follower neuron (Lateral Pyloric, LP neuron) (Fig. 2A). We identified a conductance parameter space for each cell, defined as the multidimensional space formed by the maximal conductance values of all (known and measurable) ionic currents. To assess changes in this parameter space, we used a simple metric: the total conductance variance,*i*.*e*. the sum of the variances of all measured ionic conductances in an identified cell across a population of animals. We show that the total variance, akin to the volume of the entire conductance parameter space, increases after decentralization in both PD and LP neurons. Interestingly, not all conductances increase but they change differently in the two neurons examined. We further observe that this parameter space is reduced again (*i*.*e*. is not different from control) when neuromodulators are supplied exogenously. Finally, when natural activity is artificially maintained after decentralization, the control conductance parameter space remains at control levels but in a cell-type-dependent manner. Importantly, these changes cannot clearly be attributable to any single conductance, suggesting that the cells regulate the entire parameter landscape.

## Materials and Methods

### Animals

Male Jonah crabs, *Cancer borealis*, were purchased from local fish markets in Newark, NJ, and New York City, NY. They were housed in our animal tanks at around 11–12°C in circulating, aerated artificial ocean water (Instant Ocean, Blacksburg, VA, USA). The data presented in this study were collected between April 2023 and October 2024.

### Dissection and desheathing

Before dissection, crabs were anesthetized by placing them in ice for approximately 30 min. The stomatogastric nervous system (STNS) (Fig. 1) is situated dorsally on the crab’s stomach. The STNS was then dissected as described (Gutierrez and Grashow, 2009) and pinned dorsal side up on a Sylgard-lined Petri dish.

The STNS consists of 4 ganglia: a pair of commissural ganglia (CoGs), the esophageal ganglion (OG), and the stomatogastric ganglion (STG) (Fig. 1). The sheath over the dorsal side of the STG was excised with fine tungsten pins to expose STG neuron somata for electrode impalement. The STG in *C. borealis* contains 26 neurons, including the pyloric neurons (Fig.2A). The CoGs and the OG contain the cell bodies of modulatory neurons whose axons run through the stomatogastric nerve (*stn*) (Fig. 1) and release a variety of neuromodulators in the STG neuropil (Marder et al., 2022). In the presence of neuromodulators, and at ∼12°C, the pyloric network generates a regular rhythm with a cycle frequency of ∼1 Hz (Fig. 1, 2A right, control). All our experiments were performed at ∼12°C.

### Decentralization

At the time of dissection, an oval-shaped well was first built around the STG using petroleum jelly. After identifying and impaling neuronal somata with two sharp electrodes each and waiting for ∼20 minutes for neuronal activity to stabilize, the STG was superfused with physiological saline containing 10^-6^ M tetrodotoxin (TTX, Alomone Labs). TTX blocks the voltage-gated Na^+^ current and thus the generation of action potentials in STG neurons and in neuromodulatory projection neurons. As a consequence, the release of neuromodulators from projection neurons’ axonal terminals in the STG neuropil is terminated and, thus, the STG is effectively decentralized. After action potentials were completely blocked (∼10 minutes), TTX was switched to 10^-7^ M for the remainder of the experiment.

### Solutions

Normal physiological saline: *C. borealis* saline was made of NaCl (440 mM), MgCl_2_ (26 mM), CaCl_2_ (13 mM), KCl (11 mM), Tris base (10 mM), and maleic acid (5 mM). pH was adjusted to 7.4–7.5 with either maleic acid or Tris as needed.

Mn^++^ saline: To measure *I*_Kd_, we replaced the normal Ca^++^ with 12.9 mM Mn^++^ + 0.1 mM Ca^++^. This blocks Ca^++^ channels and consequently *I*_KCa_ (Golowasch and Marder, 1992b).

Custom synthesized Proctolin (RS Synthesis, Louisville, KY, sequence RYLPT) was dissolved in distilled water and stored as 10^-3^ M aliquots at −20°C. Proctolin stock was thawed and diluted immediately before use in saline to a final concentration of 10^-6^ M (plus 10^-7^ M TTX).

### Electrophysiology

Intracellular data reported here are from Pyloric Dilator (PD) and Lateral Pyloric (LP) neurons. PD neurons are part of the pacemaker group, and the LP neuron is a follower neuron. PD and LP neurons were identified by matching their intracellular activity to their corresponding extracellular activity on the pyloric dilator (*pdn*) and lateral ventricular nerves (*lvn*), respectively (Fig. 1).

We used the two-electrode voltage clamp (TEVC) technique to measure ionic currents with intracellular solution-filled electrodes (Hooper et al., 2015) impaled in the neuronal somata. The currents were measured at 11 – 13°C using a current injection electrode with a resistance of 25 to 28 MΩ and a voltage recording electrode with a resistance of 31 to 35 MΩ. Electrodes were pulled from borosilicate capillaries with filament. Intracellular signals were amplified using an Axoclamp 2B intracellular amplifier (Molecular Devices, San Jose, CA, USA), and all recordings were digitized at 5 kHz (Digidata 1322A, Molecular Devices, San Jose, CA, USA) and recorded with Clampex 10.6 (Molecular Devices).

Voltage-clamp measurements in the control condition were initiated no less than 10 min after cessation of action potentials to ensure that Na^+^ currents were fully blocked. We impaled one PD and one LP neuron simultaneously with two microelectrodes each. When currents were measured from one of these cells, the other was clamped at −50 mV. The protocols to record each current were identical for both cells and as follows:

*I*_Leak_: A step protocol with a holding voltage, V_h_ = −40 mV, and 800 msec voltage steps were applied from −70 to −45 mV in 5 mV increments. *I*_Leak_ was measured at the end of the pulse, and a regression line was calculated across the range of voltages where the slope of the I-V curve is lowest (typically −60 to −50 mV). The slope of this curve is the leak conductance, glk.

*I*_A_ (Fig. 3B, top trace): This outward K^+^ current activates at depolarizing voltages, *i*.*e*. −30 mV and higher, completely inactivates at −40 mV, and nearly completely de-inactivates at −90 mV (Fig. 2B) (Golowasch and Marder, 1992b). Another outward K^+^ current, called Ca^++^-dependent high threshold potassium current (*I*_HTK_), also activates at the same voltages but does not inactivate. To measure *I*_A_, we first measured *I*_HTK_, with 800 msec steps from −60 to +30 mV in 10mV increments from V_h_ = −40 mV. Another set of identical voltage steps was then applied from V_h_ = −90 mV to de-inactivate *I*_A_, and the *I*_HTK_ current set was subtracted. The resulting difference is the leak-subtracted *I*_A_. To calculate gA, the peak current evoked at +20 mV and at ∼12 msec after the start of the test pulse was divided by the driving force, assuming *E*_K_ = −80mV.

**Figure 3.**
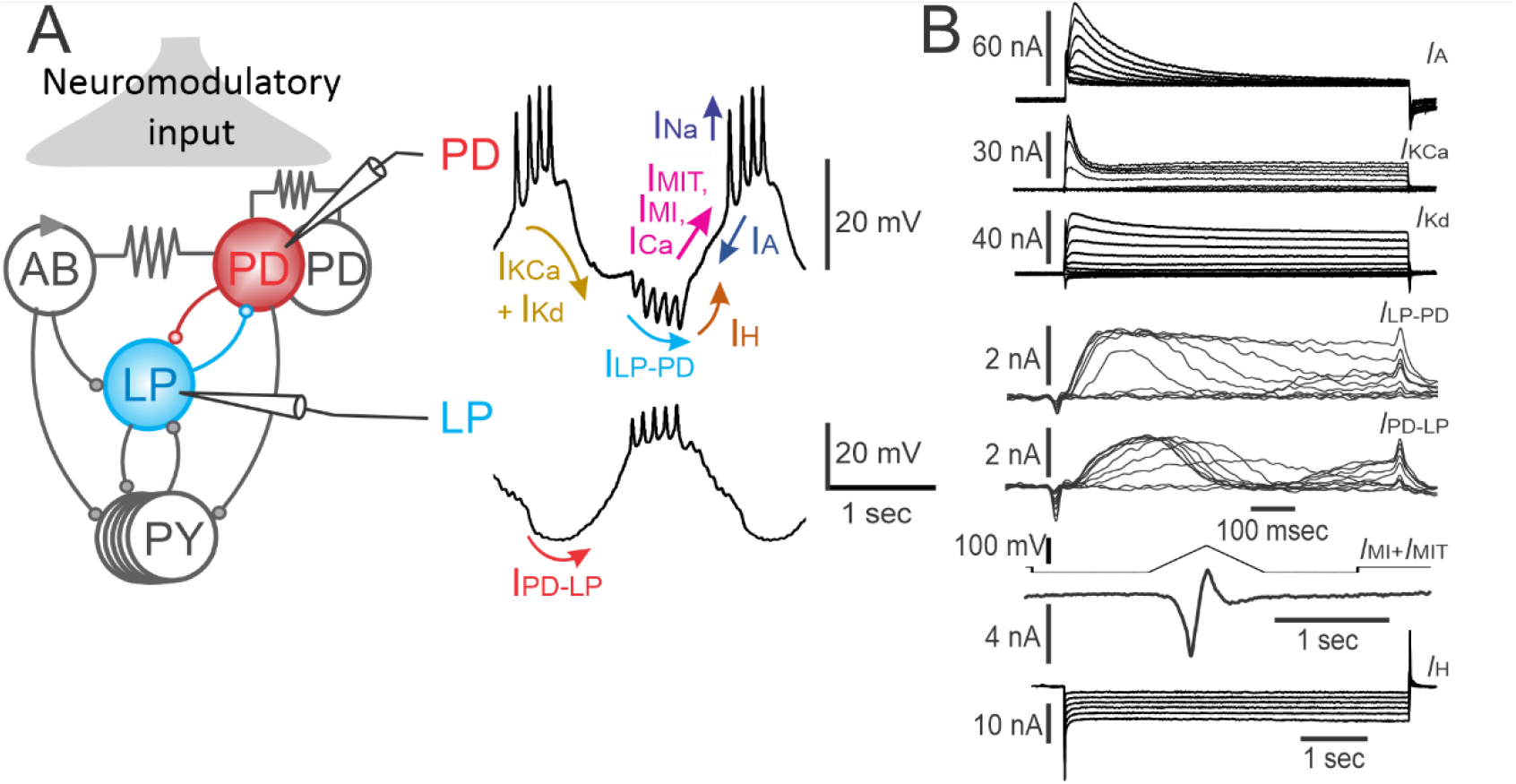
Pyloric network neurons generate regular oscillatory activity due to the interplay of several ionic currents. (**A**) Schematic diagram of pyloric network (left), highlighting the activity of the two neurons shown on the right. Ionic current labels indicate the approximate time when, and voltage range where, these currents are active during an oscillatory cycle (right). (**B**) Examples of seven of the ionic currents labeled in A measured in one of the PD neurons, and in the LP neuron (*I*_PD-LP_) using voltage and pharmacological manipulations as described in Methods. The leak current traces can be seen in the lower voltage range of the current set labeled *I*_Kd_. Voltage traces are only shown for the *I*_MI_+*I*_MIT_ current (ramps). All other currents are measured using voltage clamp steps. The time scale for the top 5 currents is the same and is shown under *I*_PD-LP_. *I*_Na_ and *I*_Ca_ cannot be measured in the same cell as the others for technical reasons (Methods) and were excluded from this study.

*I*_Kd_ (Fig. 2B, third trace): *I*_HTK_ is a current made up of two K^+^ currents, *I*_Kd_ and *I*_KCa_. To determine *I*_Kd_, we used the *I*_HTK_ protocol mentioned above but measured in Mn^++^ saline (Fig. 2B; here shown without leak subtraction). To calculate g*Kd*, the *I*_Kd_ measured at the end of the 800 msec+20 mV pulse was leak-subtracted offline using the leak current measured in Mn^++^ saline and then divided by the driving force, assuming *E*_K_ = −80 mV.

*I*_KCa_ (Fig. 3B, second trace): To determine *I*_KCa_, we used the *I*_HTK_ current protocol in normal saline and leak currents were then subtracted. The leak-subtracted *I*_Kd_ measured as described above was then subtracted (Fig. 2B). To calculate gHTK, the peak of *I*_KCa_ was measured at +20 mV and at ∼12 msec from the start of the current pulse, and divided by the driving force, assuming *E*_K_ = −80mV.

*I*_H_ (Fig. 3B, bottom trace): This current activates with hyperpolarizing pulses below −40 mV. We measured it with 4 sec long pulses to −30 to −110 mV in 10 mV increments from V_h_ = −40 mV in normal saline. To calculate gH, the maximum amplitude was measured at the end of the 4 sec pulse at −110 mV, leak subtracted, and then divided by the driving force, assuming *E*_H_ = −10 mV (Golowasch and Marder, 1992b).

*I*_MI_+*I*_MIT_ (Fig. 3B, sixth trace): These neuromodulator-activated inward currents activate together from V_H_ = −80mV in the presence of several different neuromodulators (Swensen and Marder, 2000), and, although *I*_MIT_ slowly inactivates at depolarized voltages, we did not attempt to separate them, given the complex interactions of their voltage dependencies (Schneider et al., 2021). The compound *I*_MI_+*I*_MIT_ is thus described here as *I*_MI_ and was calculated as the difference between currents measured in the presence of 10^-6^ M proctolin minus the current measured in normal saline (Golowasch and Marder, 1992a) as follows: we voltage clamped the neuron at −80 mV for 1 sec and applied a voltage ramp to +20 mV and back to −80 mV at 200 mV/sec. We repeated this five times at 10 sec intervals and averaged the currents. Between sweeps, the cell was held at V_H_ = −60 mV. gMI was calculated as the peak negative current (usually around −10 mV) divided by the driving force. For this current, we measured its equilibrium potential each time from the I-V curve, which usually lies between 0 mV and +20 mV.

*I*_Syn_ (Fig. 3B, fourth and fifth traces): To measure *I*_syn_, the postsynaptic neuron was held at −50 mV, and the presynaptic neuron was depolarized with 800 msec pulses from −60 to +20 mV in 10 mV increments from V_h_ = −90 mV. The peak synaptic current was measured at +15 mV, and gsyn was calculated by dividing this peak current by the driving force, assuming *E*_syn_=-70 mV for the LP-to-PD synapse and *E*_syn_= −80 mV for the PD-to-LP synapse (Martinez et al., 2019).

The voltages chosen for the calculation of the conductances of each current are such that they are as close as possible to their corresponding maximal conductances.

It is, unfortunately, technically not possible to measure *I*_Na_ and *I*_Ca_ in the same cells when measuring the currents listed above. *I*_Na_ is extremely fast and generated far away from the soma, where the electrodes are placed, and large capacitive artifacts plus space clamp issues preclude a decent voltage clamp measurement of this current. Likewise, *I*_Ca_ is a very small current that can only be measured after pharmacologically blocking all K^+^ currents with intracellular Cs^+^ (Graubard and Hartline, 1991; Golowasch and Marder, 1992b) and also suffers from significant space clamp errors (Graubard and Hartline, 1991; Golowasch and Marder, 1992b; Kloppenburg et al., 2000).

Data were discarded if the neuron’s input resistance dropped below 5 MΩ during the experiment and/or if the electrode offset was greater than ± 5 mV at the end of the experiment.

### Experimental sets

We performed three sets of experiments. Each experiment began with identifying PD and LP neurons, impaling both cells with two electrodes each, decentralizing the STG by bathing it in 10^-6^ M and then 10^-7^ M TTX (see above), and measuring the 7 ionic currents in each cell shortly thereafter (Control). The STG was then maintained on the electrophysiology setup at ∼12°C, continuously superfused with saline containing 10^-7^ M TTX for 8 hours, after which time all currents were measured again. The experimental sets differed based on the treatment applied during the 8 hours after decentralization (Fig. 4A).

**Figure 4.**
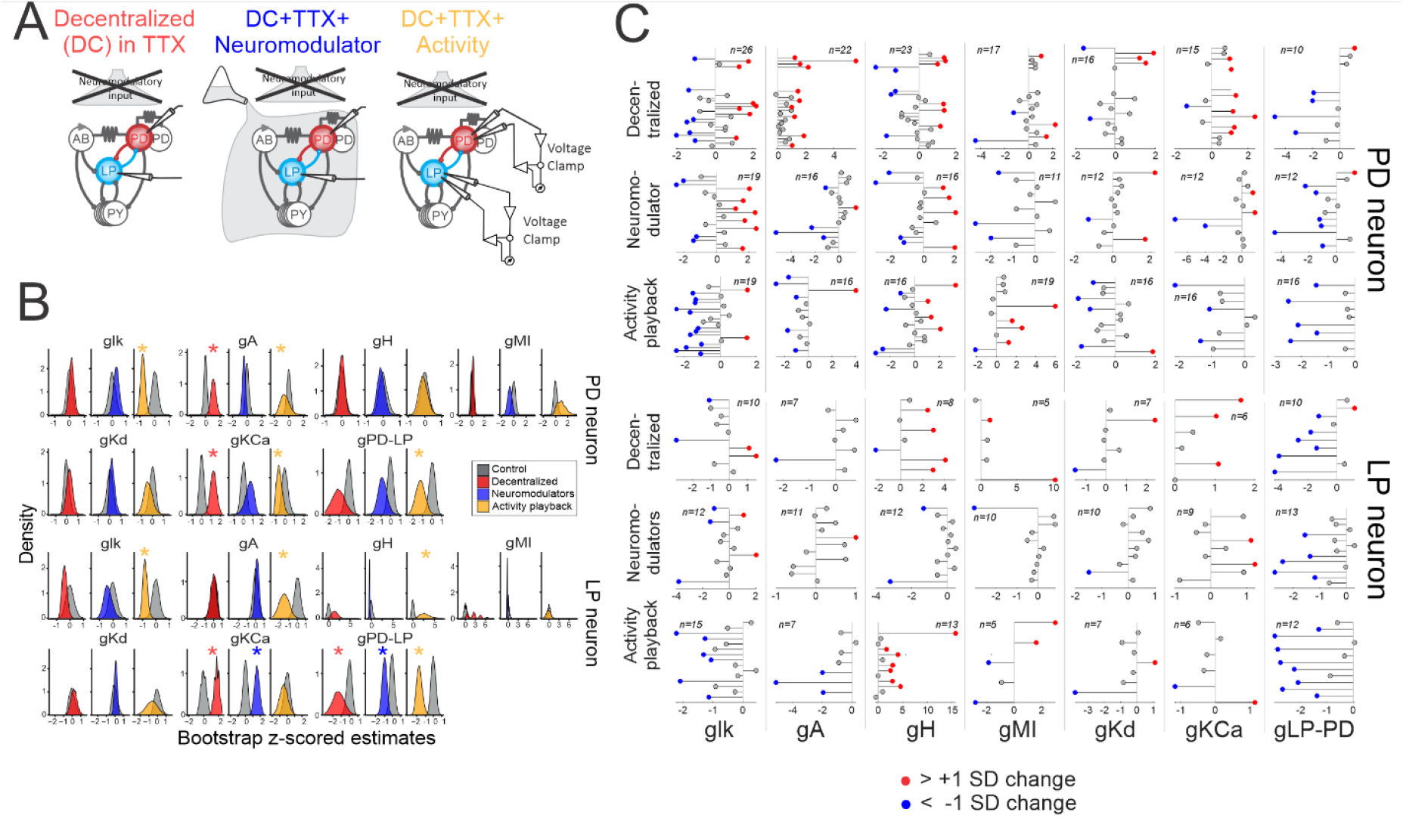
Decentralization alters conductance values and their variance. (**A**) Diagrams illustrating experimental manipulations (see main text and Methods for details). (**B**) Conductance distributions under 4 conditions: *Control* (gray), *Decentralized* (red), *Decentralized + Neuromodulator* (purple) and *Decentralized + Activity playback* (yellow). Conductances were z-scored and bootstrapped 10,000 times. All data were paired and the control (gray) distributions are specific to each cell and condition. * indicates statistical significant differences relative to their respective control data (gray) (see Table 1). (**C**) Paired conductance value (μS) changes (red, blue, and gray dots) for each recorded cell relative to its own control (beginning of line) under the 3 experimental conditions listed (A). Red dots indicate a positive change, blue dots are negative changes, and gray dots are cases where there was no change in conductance.

1. Decentralized: After the Control current measurements, the electrodes were pulled out of the cells (maintaining the impalement without injecting current through the electrodes frequently resulted in clogging of the electrodes and replacing clogged electrodes after hours of impalement invariably destroys the cells). At 8 hours, the same neurons were re-impaled, and the same set of currents was measured.
2. Decentralized + Neuromodulator: As in the first set, electrodes were pulled out of the cells after initial current measurements immediately after decentralization, and the STG was perfused with the saline containing 10^-6^ M proctolin + 10^-7^ M TTX for 8 hours. At that time, the same cells were re-impaled, proctolin was quickly washed out, and the same set of current measurements was performed.
3. Decentralized with Activity Playback: At the start of each experiment, and before TTX was applied, the natural activities of both PD and LP neurons were recorded intracellularly for 10 minutes. These recordings were saved for subsequent activity playback. After the Control current measurements were completed, both cells were kept impaled and voltage-clamped to their corresponding prerecorded intracellular voltages for 8 hours. The prerecorded voltages were trimmed and repeatedly applied without interruptions to mimic a continuous voltage trace. Then currents were measured again as before.

**Table 1.**
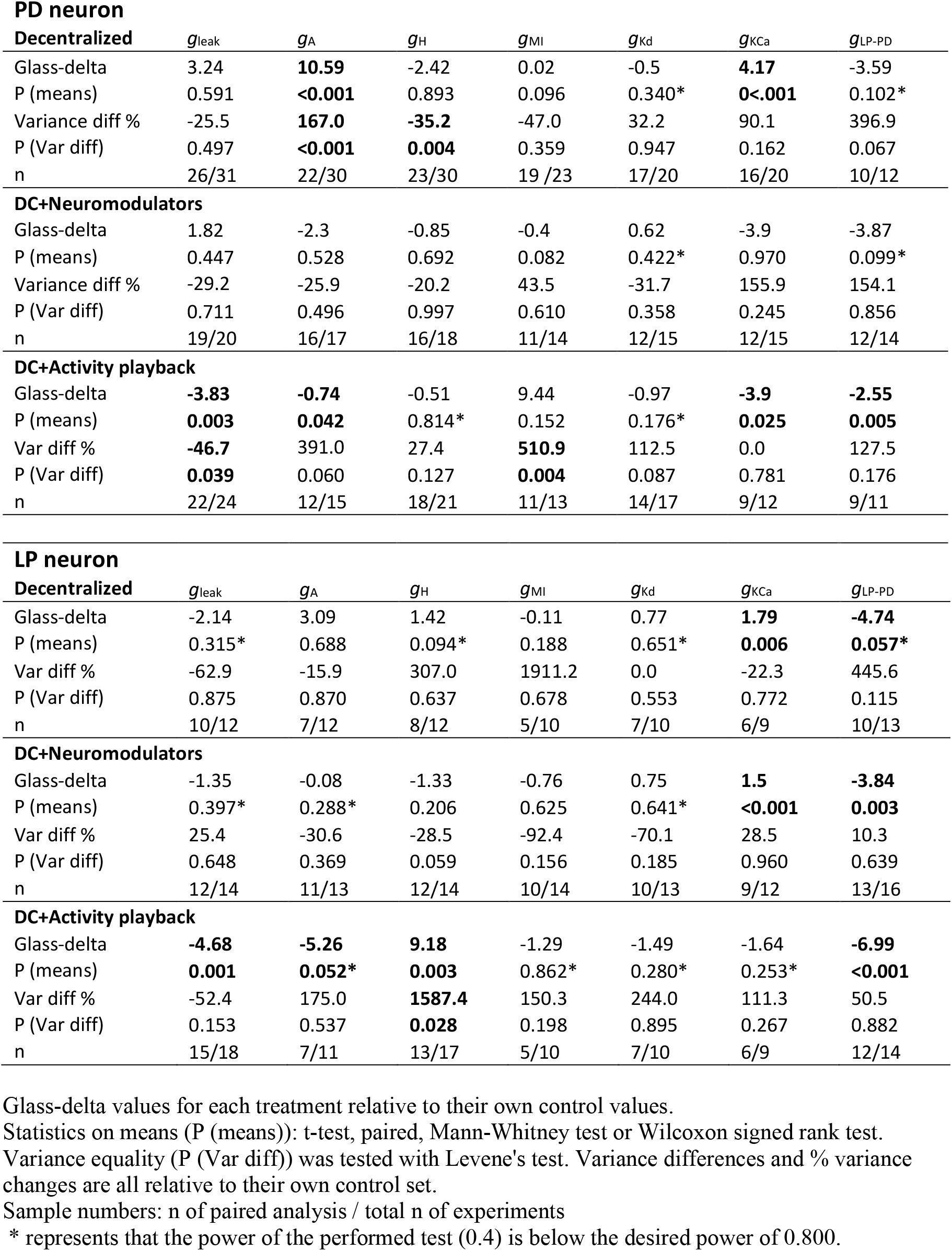
Effect size and variance changes of single conductance changes. Glass-delta values for effect size and % variance difference are reported. Bold indicates statistical significance at P<0.05.

| **PD neuron** |  |  |  |  |  |  |  |
| --- | --- | --- | --- | --- | --- | --- | --- |
| **Decentralized** | $g_{leak}$ | $g_A$ | $g_H$ | $g_{MI}$ | $g_{Kd}$ | $g_{KCa}$ | $g_{LP-PD}$ |
| Glass-delta | 3.24 | **10.59** | -2.42 | 0.02 | -0.5 | **4.17** | -3.59 |
| P (means) | 0.591 | **<0.001** | 0.893 | 0.096 | 0.340* | **0<.001** | 0.102* |
| Variance diff % | -25.5 | **167.0** | **-35.2** | -47.0 | 32.2 | 90.1 | 396.9 |
| P (Var diff) | 0.497 | **<0.001** | **0.004** | 0.359 | 0.947 | 0.162 | 0.067 |
| n | 26/31 | 22/30 | 23/30 | 19 /23 | 17/20 | 16/20 | 10/12 |
| **DC+Neuromodulators** |  |  |  |  |  |  |  |
| Glass-delta | 1.82 | -2.3 | -0.85 | -0.4 | 0.62 | -3.9 | -3.87 |
| P (means) | 0.447 | 0.528 | 0.692 | 0.082 | 0.422* | 0.970 | 0.099* |
| Variance diff % | -29.2 | -25.9 | -20.2 | 43.5 | -31.7 | 155.9 | 154.1 |
| P (Var diff) | 0.711 | 0.496 | 0.997 | 0.610 | 0.358 | 0.245 | 0.856 |
| n | 19/20 | 16/17 | 16/18 | 11/14 | 12/15 | 12/15 | 12/14 |
| **DC+Activity playback** |  |  |  |  |  |  |  |
| Glass-delta | **-3.83** | **-0.74** | -0.51 | 9.44 | -0.97 | **-3.9** | **-2.55** |
| P (means) | **0.003** | **0.042** | 0.814* | 0.152 | 0.176* | **0.025** | **0.005** |
| Var diff % | **-46.7** | 391.0 | 27.4 | **510.9** | 112.5 | 0.0 | 127.5 |
| P (Var diff) | **0.039** | 0.060 | 0.127 | **0.004** | 0.087 | 0.781 | 0.176 |
| n | 22/24 | 12/15 | 18/21 | 11/13 | 14/17 | 9/12 | 9/11 |
| **LP neuron** |  |  |  |  |  |  |  |
| **Decentralized** | $g_{leak}$ | $g_A$ | $g_H$ | $g_{MI}$ | $g_{Kd}$ | $g_{KCa}$ | $g_{LP-PD}$ |
| Glass-delta | -2.14 | 3.09 | 1.42 | -0.11 | 0.77 | **1.79** | **-4.74** |
| P (means) | 0.315* | 0.688 | 0.094* | 0.188 | 0.651* | **0.006** | **0.057\*** |
| Var diff % | -62.9 | -15.9 | 307.0 | 1911.2 | 0.0 | -22.3 | 445.6 |
| P (Var diff) | 0.875 | 0.870 | 0.637 | 0.678 | 0.553 | 0.772 | 0.115 |
| n | 10/12 | 7/12 | 8/12 | 5/10 | 7/10 | 6/9 | 10/13 |
| **DC+Neuromodulators** |  |  |  |  |  |  |  |
| Glass-delta | -1.35 | -0.08 | -1.33 | -0.76 | 0.75 | **1.5** | **-3.84** |
| P (means) | 0.397* | 0.288* | 0.206 | 0.625 | 0.641* | **<0.001** | **0.003** |
| Var diff % | 25.4 | -30.6 | -28.5 | -92.4 | -70.1 | 28.5 | 10.3 |
| P (Var diff) | 0.648 | 0.369 | 0.059 | 0.156 | 0.185 | 0.960 | 0.639 |
| n | 12/14 | 11/13 | 12/14 | 10/14 | 10/13 | 9/12 | 13/16 |
| **DC+Activity playback** |  |  |  |  |  |  |  |
| Glass-delta | **-4.68** | **-5.26** | **9.18** | -1.29 | -1.49 | -1.64 | **-6.99** |
| P (means) | **0.001** | **0.052\*** | **0.003** | 0.862* | 0.280* | 0.253* | **<0.001** |
| Var diff % | -52.4 | 175.0 | **1587.4** | 150.3 | 244.0 | 111.3 | 50.5 |
| P (Var diff) | 0.153 | 0.537 | **0.028** | 0.198 | 0.895 | 0.267 | 0.882 |
| n | 15/18 | 7/11 | 13/17 | 5/10 | 7/10 | 6/9 | 12/14 |

Glass-delta values for each treatment relative to their own control values.
Statistics on means (P (means)): t-test, paired, Mann-Whitney test or Wilcoxon signed rank test.
Variance equality (P (Var diff)) was tested with Levene's test. Variance differences and % variance changes are all relative to their own control set.
Sample numbers: n of paired analysis / total n of experiments
\* represents that the power of the performed test (0.4) is below the desired power of 0.800.

For all TEVC, the amplifier gain was set to 100 (x100 V/V).

### Data analysis and statistics

In our experiments, each of the 7 conductances measured in each neuron was first normalized by z-scoring them (z_i_ = (x_i_ - µ_x_)/σ_x_), where µ is the population average of measured current and σ is its standard deviation. Thus, the z distribution z is centered at 0 and has a standard deviation of 1.

To calculate the volume of the 7-dimensional parameter space, the variances of each of the 7 z-scored conductance values were calculated and summed to obtain the total variance, representing the 7-dimensional parameter space. The difference in total variance between the control and 8-hour treatment was computed for each experimental condition. As only a single difference value was obtained per experimental condition, deriving statistical inferences from them is not possible. To overcome this problem, we used bootstrapping (10,000 iterations with replacement) to generate the distribution of variance differences. Figure 5 shows these distributions, with a 95% bias-corrected and accelerated (BCa) bootstrap confidence interval of the distribution (DiCiccio, 1996) and shown as a colored (gray or red) rectangle. The dashed black lines correspond to the null value of the difference.

**Figure 5.**
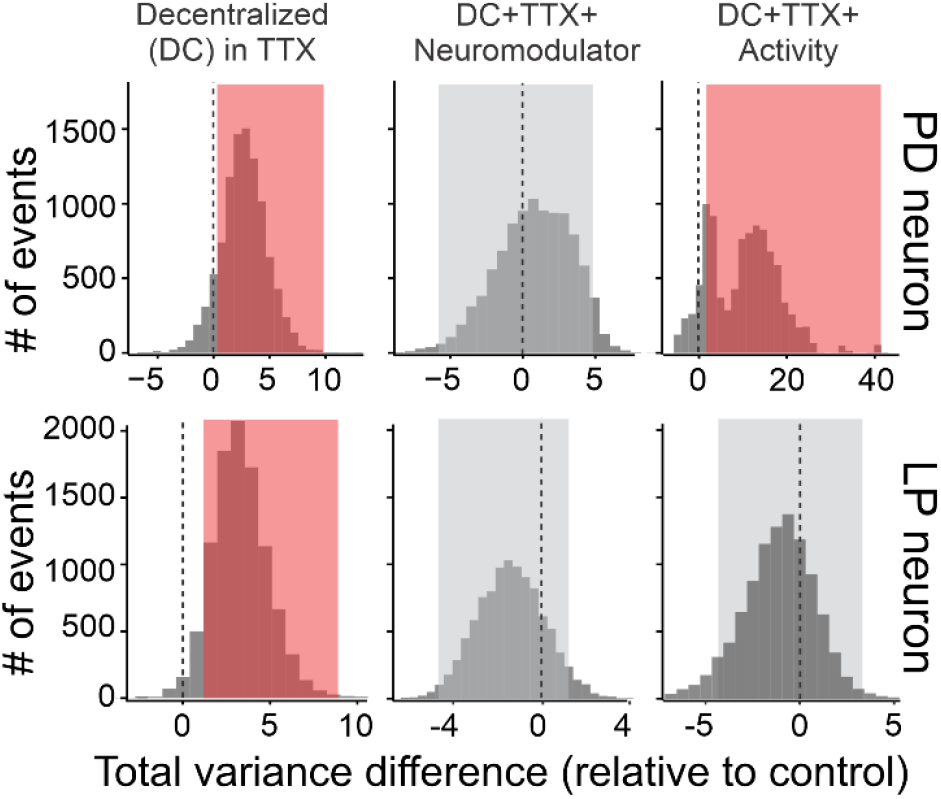
Decentralization modifies the ionic conductance space of both PD and LP neurons. The seven-dimensional conductance parameter space size (or volume) of the PD neuron (Hamood et al.) and the LP neuron (bottom) are represented by the bootstrapped difference in total conductance variance, calculated as the sum of the variances of seven z-scored and control-subtracted ionic conductances measured in each cell (Methods). For each treatment, the conductances were measured at 2 time points: shortly after decentralization and eight hours after decentralization under each of three treatments: *Decentralized* only, *Decentralized + Neuromodulator*, and *Decentralized with Activity Playback*. Black dashed lines show the null values for each distribution, indicating that there is no change in the total variance after 8 hours of treatment. Colored rectangles represent the 95% bias-corrected and accelerated (BCa) confidence interval (CI) range for the distribution of control-subtracted variance differences. Red rectangles indicate a significant departure from the null value (dashed line is outside the CI box); gray rectangles show a lack of statistical significance.

Statistical analyses were performed using RStudio and SigmaStat software packages (SPSS, Chicago, IL, USA).

## Results

### Intrinsic and synaptic currents coordinately generate the pyloric rhythm in the presence of neuromodulators

The pyloric network (Fig. 2, 3A left) has been extensively studied as a model system to understand the effects of neuromodulators and activity at the single identified neuron, synapse, and network output levels. In the presence of endogenous neuromodulators, pyloric neurons typically generate and maintain ongoing oscillatory activity (Marder and Bucher, 2007) (Fig. 1) resulting from the sequential voltage- and time-dependent activation and inactivation of multiple ionic currents (Fig. 3A, right). This is thought to be driven by a pacemaker current, the neuromodulator-activated inward current, *I*_MI_ (Bose et al., 2014) acting as a pacemaker current. However, several ionic currents profoundly affect this rhythm as illustrated by blocking different conductances of the pacemaker neurons of the network (Fig 2B). This suggests the possible diversity of pacemaking mechanisms in the network. Several of these currents have been characterized in pyloric neurons (Graubard and Hartline, 1991; Golowasch and Marder, 1992b; Schneider et al., 2021), and examples of their recordings are shown for a PD neuron in Figure 3B.

The PD neuron hyperpolarizes upon receiving synaptic inhibition (*I*_LP-PD_) from its network partner, the LP neuron, as shown in Figure 3A (right) and rebounds from this inhibition mediated in part by the hyperpolarization-activated inward current, *I*_H_. The ensuing depolarization leads to subsequent activation of a putative low-threshold calcium currents (Zhao et al., 2011) and neuromodulator-dependent inward currents, *I*_MI_ and *I*_MIT_ (Golowasch and Marder, 1992a; Schneider et al., 2021), which further depolarize the cell (Swensen and Marder, 2000; Bose et al., 2014) (here we measure these 2 currents as one and thus refer to them as *I*_MI_, see Methods).

This depolarization activates the voltage-dependent transient potassium current, *I*_A,_ with the interplay between *I*_H_ and *I*_A_ determining the rebound delay or phase and the initial spiking rate of the burst (Tierney and Harris-Warrick, 1992; MacLean et al., 2005). As the cell depolarizes, voltage-dependent sodium, *I*_Na_, as well as high-threshold calcium currents, *I*_Ca_ (Golowasch and Marder, 1992b), generate a burst of action potentials. During this depolarized phase, a calcium-dependent potassium current, *I*_KCa_, and a delayed rectifier potassium current, *I*_Kd_, are activated, repolarizing the membrane (Graubard and Hartline, 1991; Golowasch and Marder, 1992b). At this point, the PD neuron receives the next inhibitory input from the LP neuron, and the cycle repeats (Fig. 3A, right). A similar sequence of events takes place in the LP neuron.

Representative raw traces of most of these currents are shown in Figure 3B. In this study, we measured seven of these ionic currents (*I*_A_, *I*_Kd_, *I*_KCa_, *I*_H_, *I*_MI_, a voltage-independent leak current, *I*_leak_, and a synaptic current: *I*_LP-PD_ in PD and *I*_PD-LP_ in LP neurons) using various voltage and pharmacological manipulations (Methods) to build a conductance parameter space for each cell. Currents *I*_Na_ and *I*_Ca_ cannot be measured in the same cell with the other currents shown in Figure 3B, for technical reasons as explained in Methods. We simultaneously modified the neuromodulatory environment and neuronal activity of one of the two PD neurons and the LP neuron in each preparation (Fig. 4A) to assess how the conductance parameter space is impacted. However, we then independently assessed the contributions of neuromodulators and activity to the changes observed.

### Decentralization alters mean conductance values and broadens the conductance distributions

Decentralization not only blocks neuromodulatory inputs to the pyloric neurons but also alters their activities (Fig. 2A) (Luther et al., 2003; Hamood et al., 2015; More-Potdar and Golowasch, 2023). Previous studies have identified the presence of activity-dependent regulatory mechanisms that control the expression of ionic currents in pyloric neurons (Turrigiano et al., 1994; Golowasch et al., 1999; Haedo and Golowasch, 2006; Santin and Schulz, 2019).

Therefore, it is possible that observed changes in currents following decentralization may be the consequence of changes in activity rather than solely from the loss of neuromodulatory input. To evaluate the long-term effects of neuromodulators on the conductances of pyloric neurons, we carefully isolated activity-dependent effects on the ionic currents of PD and LP neurons from neuromodulator-dependent effects by using three sets of experimental conditions (Fig. 4A): i) *Decentralized*, ii) *Decentralized + Neuromodulator*, iii) *Decentralized + Activity Playback. Decentralized* preparations were maintained in saline containing 10^-7^ M tetrodotoxin (TTX) for 8 hours. TTX blocks action potential transmission along axons of neuromodulatory projection neurons, thereby preventing the release of neuromodulators in the STG neuropil. It also blocks oscillatory/bursting activity (Golowasch and Marder, 1992b). *Decentralized + Neuromodulator* preparations were maintained in normal saline containing 10^-7^ M TTX + 10^-6^ M proctolin for 8 hours. In this condition, the neurons remain inactive, and are thus deprived of activity, but not of neuromodulators. Proctolin is a neuropeptide that activates *I*_MI_ (as does pilocarpine, see Fig. 2B) in both PD and LP neurons (Golowasch and Marder, 1992a; Swensen and Marder, 2000, 2001) and can activate oscillatory activity in quiescent pyloric networks (Hooper and Marder, 1984; Marder et al., 1986) and strengthen weak activity (*e*.*g*. Fig. 2B). Proctolin also activates *I*_MIT_, which regulates burst onset in a frequency-dependent manner (Schneider et al., 2021). Here we refer to the sum of *I*_MI_ and *I*_MIT_ as *I*_MI_ (and *g*_MI_ as their total conductance). *Decentralized + Activity Playback* preparations were maintained in 10^-7^ M TTX, and the natural activities of the same cells (PD and LP neurons) prerecorded at the beginning of the experiments were continuously played back to each cell in voltage clamp for 8 hours. In this condition, the neurons are deprived of neuromodulators but not activity. The ionic currents described above (Fig. 3B) were measured immediately after TTX-mediated decentralization (and not before, because current measurements when neuronal voltage is oscillating can result in considerable measurement errors) and constitute control measurements (Methods). At such an early time post-decentralization, we believe that the long-term effects of decentralization are not likely to have significantly occurred. The next measurement was taken 8 hours after decentralization for each of the three manipulations described above (Fig. 4A). Maximal conductances were calculated at specific voltages as described in Methods.

Our results revealed large and significant changes in the mean values of most conductances in both neuron types across the three experimental conditions relative to control (Fig. 3B, Table 1). In PD neurons, two currents significantly changed their conductance in the *Decentralized* condition: *g*_A_ and *g*_KCa_, both of which showed a very large increase (effect size>>1), consistent with what we have previously observed (Khorkova and Golowasch, 2007). None of the other conductances showed a significant change. However, *g*_LP-PD_ showed a large decrease (effect sizes<1) under the *Decentralized* condition relative to *Control*, consistent with results obtained in lobsters (Thoby-Brisson and Simmers, 2002), and *g*_H_ and *g*_leak_ also showed large (but not significant) changes (effect sizes > ±1).

In LP neurons, we observed different changes: *g*_KCa_ and *g*_PD-LP_ showed large and significant changes after decentralization (effect sizes > ±1), while *g*_lk,_ *g*_A_, and *g*_H_ showed relatively large (but not significant) changes (effect sizes > ±1). Although we were not able to measure calcium currents due to the lack of effective pharmacology to isolate them (see Methods), the strong increase of *g*_KCa_ in both PD and LP neurons suggests possible underlying changes in calcium currents in addition to (or perhaps instead of) changes in *g*_KCa_. As *I*_MI_ is directly activated by neuromodulators and is thought to be the pacemaker current in the pyloric network, we expected to observe a considerable compensatory (homeostatic) increase in *g*_MI_ after prolonged decentralization, as has been observed before in LP neurons for a different neuromodulator (Lett et al., 2017). Interestingly, we did not detect any notable change in *g*_MI_ in either PD or LP neurons across the conditions tested. These observations raise the possibility that *I*_MI_ may not be the sole pacemaker current in pyloric neurons, and that there could be additional neuromodulator-independent pacemaking mechanisms that can become activated under certain conditions.

As expected, in the presence of proctolin in the *Decentralized + Neuromodulator* condition, the neuromodulator restored control levels of all voltage-dependent conductances (Khorkova and Golowasch, 2007), and one synaptic current (*g*_LP-PD_), as can be seen in Figure 4B and Table 1, suggesting that one neuromodulator added during the 8 hours post-decentralization period can prevent most of the changes induced by decentralization. In all LP neuron conductances and 6 of 7 PD neuron conductances, the distributions of the conductances in the presence of proctolin (Purple) sit closer to their *Control* distributions than the decentralized distributions (Red), as can be seen from the decrease of the absolute value of the effect size of each current. However, the synaptic conductance of PD-to-LP (*g*_PD-LP_) showed a significant decrease in the *Decentralized + Neuromodulator* condition relative to *Control* values in the presence of bath-applied proctolin. In the short term, proctolin has been shown to influence synaptic strength and dynamics by modulating both the graded- and the spike-mediated components of the LP to PD synapse (Zhao et al., 2011). However, until now, there has been no evidence of long-term effects of neuromodulators on crab pyloric synapses.

To assess the effect of activity on the changes induced by decentralization, we played back their own pre-decentralization activity on both PD and LP neurons (see Methods). Several conductance distributions shifted from *Decentralized* toward *Control* levels after 8 hours of *Decentralized + Activity playback* (Table 1, Fig. 4B). However, some (*g*_leak_, *g*_A_, *g*_KCa_, and *g*_LP-PD_ in PD neurons, and *g*_leak_, *g*_A_, *g*_H_, and *g*_PD-LP_ in LP neurons) showed significant changes relative to their control levels (Fig. 4B, compare yellow to gray distributions) sometimes with large effect sizes. While some of these currents underwent a restoration towards control levels (*g*_A_, *g*_KCa_ and *g*_LP-PD_ in PD neurons), surprisingly, all others shifted even further after *Activity playback* than in the *Decentralized* condition. The changes described above are also clearly observed in most individual experiments (Fig. 4C).

Overall, our results strongly suggest that decentralization induces a membrane restructuring not only at the level of individual conductances but across the entire conductance landscape. Importantly, in every experimental condition, the conductance distributions broadened relative to their control distribution in both cell types.

A second important observation revealed in Figure 4 and Table 1 is the very large and significant increase in the variance of *g*_A_ and a much smaller but, also significant, decrease in the variance of *g*_H_ upon decentralization. Additionally, 3 other PD neuron conductances showed large increases in variance and 3 much smaller decreases; however, none were statistically significant. Overall, the total conductance variance of the PD neuron increased by 578.5%. At the same time, the LP neuron had a large total variance increase of 2562.7%, but none of the individual conductance variance changes was statistically significant (Table 1, top panels). All variance changes were fully reversed after restoring neuromodulatory input, without any of the individual conductances showing a statistically significant difference from control, with the total variance remaining high (246.5%) in PD neurons, but reduced (−157.4%) in LP neurons (Table 1, middle panels). The picture is more confusing when restoration of only activity is considered, with the variance of PD neurons increasing significantly for *g*_MI_ and decreasing significantly for *g*_leak_ (overall 1122.6%), and the variance of LP neurons significantly increasing for *g*_H_ (overall 2266.1%) (Table 1, bottom panels).

### Decentralization alters the conductance parameter space of both PD and LP neurons

As described above, the results obtained at the single conductance level (Fig. 4) strongly suggest that conductance distribution (a proxy for variance) increases with decentralization, and when the neuromodulatory environment is maintained, the conductance distributions remain mostly unchanged relative to control distributions (although the effect of activity playback is less clear). The observed increases in overall variance are consistent with our hypothesis that neuromodulators (and/or activity) constrain the entire conductance parameter space of STG pyloric neurons and, conversely, decentralization increases it. To more rigorously test the hypothesis that the entire conductance parameter space is regulated by neuromodulators in these cells, we computed the total variance of the 7 measured ionic conductances expressed by PD and LP neurons by simply summing the variances of all seven individual conductances. However, prior to this, we z-scored each conductance to ensure they are all on a comparable scale (Methods) and then measured the difference between total variances observed at 0 hours (*Control*) and 8 hours later, after treatment with one of the three conditions described before (Fig. 4A). This difference quantifies the change in the conductance parameter space in response to the presence or absence of neuromodulators and/or activity. A positive difference indicates expansion of the parameter space; a negative difference suggests contraction. To enable a rigorous statistical comparison, we applied a bootstrap resampling procedure (10,000 iterations). At each iteration, random samples with replacement were drawn in parallel for all 7 conductances in two groups: one from the *Control* dataset and the other from the 8-hour treatment dataset. Experiments in these two datasets were paired. The total variances were then computed for each case, and the difference was calculated. Fig. 5 presents the resulting frequency distributions of variance differences for each cell type. The colored (red and gray) rectangles denote the 95% confidence intervals (bias-corrected and accelerated CIs (DiCiccio, 1996)), while the black dashed lines represent the null values for each treatment (no variance difference). If the null value falls outside the CI for each treatment, it would indicate that a significant shift in the variance distribution.

Our results show that the difference in total variances between the *Control* and *Decentralized* conditions significantly increases in both PD and LP neurons (Fig. 5, left). We take this as an indication that the conductance parameter spaces in both PD and LP neurons expanded significantly after decentralization. When a neuromodulator (proctolin) is present for eight hours in the *Decentralized + Neuromodulator* condition, the null value falls within the CI range for both PD and LP neurons, indicating that the presence of the neuromodulator prevents the expansion of parameter space in both cells (Fig. 5, center), consistent with our hypothesis that neuromodulators constrain the conductance parameter space. Lastly, our results also show that activity alone (8 hours of *Decentralized + Activity playback* in the absence of neuromodulators) can also prevent the expansion of the conductance parameter space, but this appears to be a cell-type-dependent effect: in PD neurons, the presence of activity in a decentralized condition does not reduce the global variance induced by decentralization, in contrast to LP neurons, where it does (Fig. 5, right).

All together, our results strongly support the hypothesis that decentralization induces membrane restructuring across the entire conductance parameter space.

## Discussion

Here, we provide evidence that neuromodulators in the STG pyloric network of crabs reduce the size of the conductance parameter space of two central neurons of the network: a member of the pacemaker kernel, the PD neuron, and the only neuron that provides feedback to the pacemaker of the network, the LP neuron. We show that, after decentralization, about half of the individual ionic currents change their conductances in two ways: the means tend to increase and the conductance distributions in a population of cells broaden (the variance increases). When (only) neuromodulators are restored, the changes tend to shift back towards control levels in both cells. When only activity is restored, the results are less clear but there is a tendency for the distributions to shift back towards control levels as well (Fig. 4). These data are consistent with previously observed changes induced by decentralization (Khorkova and Golowasch, 2007; Santin and Schulz, 2019) that showed a reduction in the strength of correlated expression of pairs of ion channels in crab PD neurons, which corresponds to an increase in total variance of the correlated distributions. In 2007, Khorkova and Golowasch showed that proctolin can restore the conductance correlations, and in 2019 Santin and Schulz, using single-cell mRNA expression data, showed that activity can also restore a number of these correlations. However, neither of these two studies clearly shows the changes in the size of the conductance space. In our study, we sampled a large amount of data (especially for PD neurons) to assess potential changes in the width of the distributions and measured the global conductance variance of each cell as an estimate of the volume of their parameter space to disambiguate some of the less clear changes observed at the single conductance level. We observe a clear and significant increase in the global variance of both cells after decentralization, which is restored by neuromodulators and (in part) by activity.

We conclude that neuromodulators and activity regulate the size (volume) of the global parameter space of both pyloric neurons, as hypothesized (Golowasch, 2019a). Given that activity is spontaneously recovered after a prolonged decentralization even in the absence of neuromodulators (Fig. 2A), we propose that these conductance changes constitute a mechanism by which the system finds a new set of conductance values that allows the reemergence of oscillatory activity. Furthermore, this emergence of new sets of conductance values that support recovery of the oscillatory activity implies a possible shift in the rhythm (pacemaking) generating mechanism as a consequence of the perturbations to the network (*e*.*g*. loss of neuromodulation). A similar shift has been shown to occur during development in mammalian substantia nigra neuronal bursting (Chan et al., 2007) and in response to the removal of neuromodulators in the mammalian respiratory system (Telgkamp et al., 2002). Thus, this ability of rhythm-generating networks to reconfigure their rhythm-generating mechanisms is likely to be a general one. This ability may increase the effective complexity and consequently the robustness of the networks (Golowasch, 2023). Interestingly, if rhythm-generating mechanisms inhabit a dynamic, perhaps constantly changing, parameter space of a neuron, then studies focusing on these mechanisms should not expect to find unique mechanisms but, rather, degenerate ones instead. Our data provide a possible explanation for why we see such a large range of responses to decentralization, with rhythmic pyloric activity sometimes completely shutting down and sometimes being affected only mildly (compare the effects of decentralization in 2 different animals, shown in Figures 2A and 2B). Possible mechanisms include a Ca^++^ current and a Ca^++^-dependent K^+^ current based oscillator (Daur et al., 2016) and a neuromodulator-activated current based mechanism (Bose et al., 2014), or combinations of parameters of these three currents, which may produce a continuum of potential mechanisms.

We observe differences in the role that neuromodulators and neuronal activity play in constraining the conductance parameter space between different cell types. Our results suggest that there could be neuromodulator-dependent and activity-dependent intracellular pathways operating in parallel (Golowasch, 2019b), homeostatically regulating not just ion channel expression levels but also maintaining the constrained structure of the entire conductance parameter space. In some cases, such as in LP neurons, these pathways may converge, as we observe both neuromodulators and activity-restoring control conditions after decentralization, whereas in other cases, such as in PD neurons, these regulatory pathways may function more independently, at least at the protein (*i*.*e*. conductance) level. It should also be noted that it is likely that the regulatory mechanisms of the transcriptional parameter space defined by the mRNA species that code for the ion channels expressed by these neurons are not the same as the regulatory mechanisms that determine the conductance parameter space, since these two parameter spaces respond differently to the same manipulations (Temporal et al., 2012).

What might be the regulatory mechanisms that control ion channel expression over long time scales, such as those observed here? Neuromodulators, through GPCR signaling, may activate second messenger systems that act as sensors of neuromodulators and activity. Some of these second messengers may also respond to activity itself, thereby activating transcription factors that regulate ion channel expression (Sugo et al., 2025). At the moment, we do not know which transcription regulatory mechanisms are responsible for these changes. This, together with the biophysical identity of the rhythm-generating mechanisms, should be fruitful future research paths.

